# Feasibility of laminar functional quantitative susceptibility mapping

**DOI:** 10.1101/2024.09.16.613070

**Authors:** Sina Straub, Xiangzhi Zhou, Shengzhen Tao, Erin M. Westerhold, Jin Jin, Erik H. Middlebrooks

## Abstract

Layer fMRI is an increasingly utilized technique that provides insights into the laminar organization of brain activity. However, both blood-oxygen-level-dependent (BOLD) fMRI and vascular space occupancy data (VASO) have certain limitations, such as bias towards larger cortical veins in BOLD fMRI and high specific absorption rate in VASO. This study aims to explore the feasibility of whole-brain laminar functional quantitative susceptibility mapping (fQSM) compared to laminar BOLD fMRI and VASO at ultra-high field. Data were acquired using 3D EPI techniques. Complex data were denoised with NORDIC and susceptibility maps were computed using 3D path-based unwrapping, the variable-kernel sophisticated harmonic artifact reduction as well as the streaking artifact reduction for QSM algorithms. To assess layer-specific activation, twenty layers were segmented in the somatosensory and motor cortices, obtained from a finger tapping paradigm, and further averaged into six anatomical cortical layers. The magnitude of signal change and z-scores were compared across layers for each technique. fQSM showed the largest activation-dependent mean susceptibility decrease in Layers II/III in M1 and Layers I/ II in S1 with up to -1.3 ppb while BOLD fMRI showed the strongest mean signal increase in Layer I. Our data suggest that fQSM demonstrates less bias towards activation in superficial layers compared to BOLD fMRI. Moreover, activation-based susceptibility change was comparable to VASO data. Studying whole-brain, layer-dependent activation with submillimeter fQSM is feasible, and reduces bias towards venous drainage effects on the cortical surface compared to BOLD fMRI, thereby enabling better localization of laminar activation.

## 1. Introduction

The rapidly expanding field of laminar functional magnetic resonance imaging (fMRI) is dedicated to evaluating functional activation in individual cortical layers (1). The cortex is comprised of six principal layers, with varying thickness depending on the specific cortical region of interest, ranging approximately from 0.1-1 mm (2). Recent advancements in fMRI technology, particularly the utilization of ultra-high field (UHF) (magnetic field strength ≥ 7 T) have facilitated fMRI with exceptional temporal and spatial resolution, reaching up to 0.35 mm isotropic resolution (3,4). However, the ability of functional imaging techniques to assess activation in specific layers also relies on the choice of contrast mechanisms. While gradient-echo blood-oxygen-level-dependent (BOLD) fMRI is highly sensitive to functional activation, it is also biased towards venous drainage effects on the cortical surface, limiting its spatial selectivity (5). On the other hand, techniques based on cerebral blood volume (CBV) and cerebral blood flow (CBF) have demonstrated the ability to better localize activation within the corresponding cortical layers (6). Vascular space occupancy (VASO) has gained popularity as a technique for assessing laminar activation, as it minimizes the impact of venous drainage effects (4-7).

Quantitative susceptibility mapping (QSM) can quantify the change in magnetic susceptibility related to functional activation, which forms the basis for the contrast observed in BOLD fMRI. Functional QSM (fQSM) has been explored in previous studies (8-12) utilizing 2D EPI with up to 1 mm isotropic resolution. These initial investigations have shown that fQSM offers improved localization capability of brain activation in cortical gray matter, making it an intriguing addition to magnitude-based BOLD fMRI (13). The objective of this study is to assess the feasibility of laminar fQSM using a 3D EPI-fQSM approach (14) versus traditional laminar BOLD fMRI and VASO.

## 2. Methods

### 2.1 Data acquisition

In accordance with Institutional Review Board approval, data were acquired in thirteen healthy volunteers after written informed consent was obtained. One subject was included in two different sessions. MRI was performed under the investigational parallel transmit system on a 7T scanner (MAGNETOM Terra, Siemens Healthineers, Erlangen, Germany) using an investigational 8Tx-32Rx head coil (Nova Medical Inc., Wilmington, MA, USA). A segmented, multi-shot gradient-echo 3D EPI research sequence (15) was employed, including skipped-CAIPIRINHA acceleration (16), navigator echoes acquired per excitation, and imaging echoes from the center of the k-space to estimate the zeroth- and first-order phase correction (17). Additionally, VASO data were acquired in four subjects using a research sequence (16,18). The 3D EPI sequence was acquired in a strictly axial orientation with left-right and head-foot phase encoding directions. The VASO sequence was acquired in strictly axial orientation with anterior-posterior and head-foot phase encoding directions, inversion times TI_1_ = 993 ms and TI_2_ = 1859 ms, four magnetic preparations per shot, and fat suppression. All other sequence parameters are listed in Table 1.

**Table 1.**
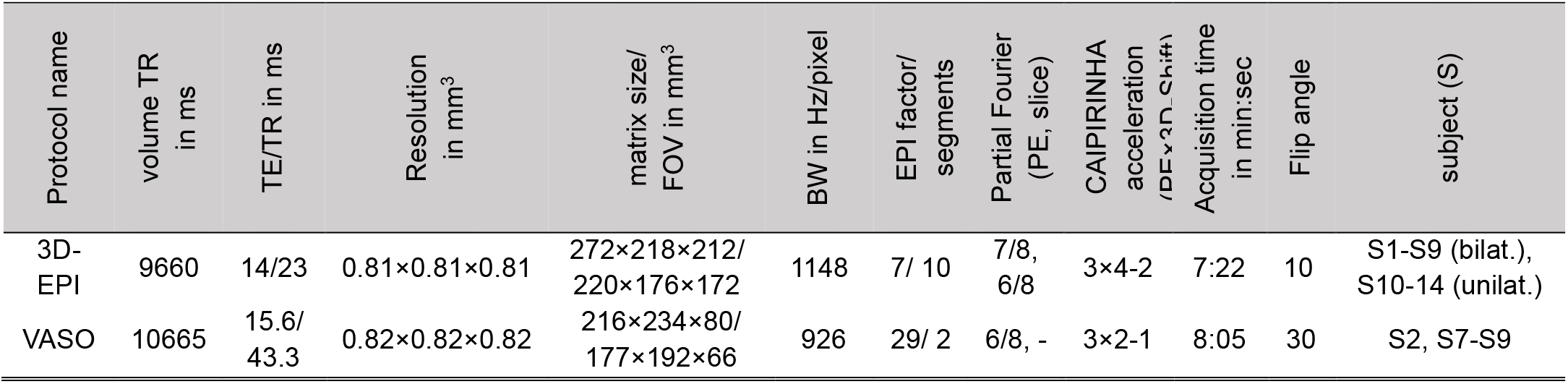
Sequence parameters and functional paradigms.

### 2.2 Stimuli and paradigms

A finger-tapping paradigm (unilateral right hand in five subjects and bilateral in nine subjects; all fingers) was performed consisting of alternating blocks of the active task and rest. Seven blocks of the active task and eight blocks of rest were obtained with three measurements acquired during each block. Each total fMRI run consisted of 45 measurements. The start and stop commands for the paradigm were displayed on an LCD monitor (NordicNeuroLab Inc. Milwaukee, WI, USA) and viewed through a mirror mounted to the head coil. The paradigms and subjects are listed in the last column of Table 1.

### 2.3 Data pre-processing and computation of functional susceptibility maps

Complex data were denoised using NOise reduction with DIstribution Corrected (NORDIC) principal component analysis (19). Susceptibility maps were computed from the phase data of each volume. The algorithm implementations in the SEPIA toolbox (20) and STISuite were used. Phase was unwrapped using 3D path-based unwrapping (21). Background field removal was performed with variable-kernel sophisticated harmonic artifact reduction for phase data (V-SHARP) using default parameters (22) and fourth-order polynomial fitting to correct for radiofrequency transmit-phase offset (20). Dipole inversion was computed using the streaking artifact reduction for QSM (STAR-QSM) algorithm (23) with default parameters. HD-BET (24) was run on CPU in ‘accurate’ mode for the average magnitude data of all volumes of each dataset to generate the brain mask used for background field removal and susceptibility map computation. This mask was further eroded using a spherical structuring element with radius of one voxel, and each volume was referenced to the mean susceptibility value of the whole brain.

### 2.4 Functional processing of the time series

For all acquisitions, the volumes were aligned to the first measurement of the time series using FSL FLIRT (25,26). For the one subject that was included for both bi- and unilateral finger tapping in two separate sessions, data were also coregistered. Quality metrics (skew, kurtosis, auto-correlation and global signal correlation) were computed for the motion-corrected time-series data using the LN_SKEW function in laynii (27).

VASO data were then co-registered and time-interpolated in Matlab. The first three measurements were discarded from each time series and a general linear model was fit including eight drift regressors to account for low frequency confounds using the spm_filter function implemented in SPM12 (28). Fitted low frequency drift was subtracted from the fQSM, BOLD fMRI and VASO data. BOLD correction was performed by dividing the BOLD-contaminated VASO signal by the BOLD signal as described in Huber et al. (27,29). Maps of susceptibility change, percentage of BOLD and VASO signal change, respectively, as well as z-scores were computed from the corrected time series data. All data were up-sampled by a factor of four and separate masks were manually drawn on three adjacent slices in the motor (M1) and somatosensory cortex (S1) on fQSM and VASO data, respectively, using FSLEyes. The mask for M1 was delineated according to the border of BA4a/BA4p (30) and the folding location of the lateral end of the hand knob as described in Huber et al. (5). The mask for S1 was drawn in the S1 cortex directly posterior to the M1 mask. These masks were used to segment 20 equidistant layers with the LN_GROW_LAYERS function in laynii (27). For the remainder of this manuscript, according to Wagstyl et al. (2), it is assumed that for M1 (S1), the segmented Layers 1 and 2 (1-2) approximately correspond to cortical Layer I, 3-5 (3-6) to Layer II, 6-9 (7-10) to Layer III, 10 and 11 (11-14) to Layer Va, 12-15 (15-16) to Layer Vb, and 16-20 (17-20) to Layer VI. Furthermore, LN_LAYER_SMOOTH was used to smooth maps of susceptibility and percentage BOLD signal change as well as z-score maps within layers with a full width at half maximum set to 1 mm. To further analyze the time series data, data were averaged across subjects, and time-averaged active and rest periods were computed (three data points each) from the 42 measurements. Cortical depth profiles were also computed by averaging all time-points from active periods. Wilcoxon rank sum test was used to assess statistical differences between the times series of subjects who performed unilateral and bilateral finger-tapping, respectively, as well as between the z-scores of fQSM, BOLD fMRI and VASO. A p-value of p < 0.05 was considered statistically significant.

## 3. Results

Figure 1 shows three axial slices of one time point as well as of the time-averaged, raw (i.e. before drift correction was applied) fQSM and BOLD fMRI data of one subject and higher order quality metrics. For mean fQSM and single-time-point fQSM few artifacts can be observed in the vicinity of large vessels which were more apparent in the skew and kurtosis maps. Auto-correlation maps as well as correlation maps which the global signal appeared relatively flat for both fQSM and fMRI.

**Figure 1.**
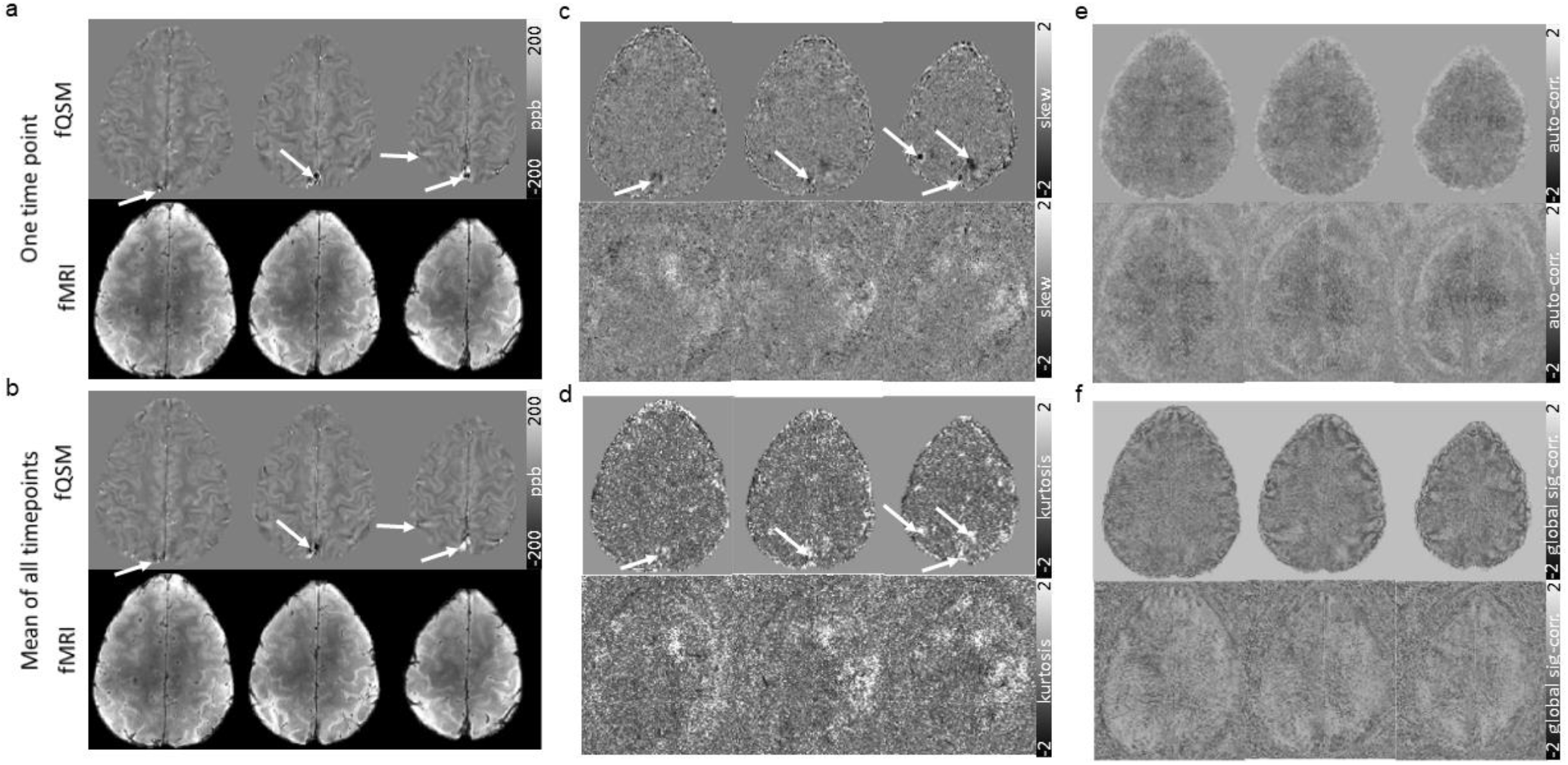
Three slices of fQSM and fMRI are shown for one example subject and one time point (a) as well as a mean image over all time points (b). Additionally, temporal skew (c), kurtosis (d), auto-correlation (e) and correlation maps with the global signal (f) are displayed. White arrows point at artifacts in fQSM in the vicinity of larger venenous vessels.

Figure 2 shows the group-averaged fQSM and fMRI data acquired from M1 and S1 (both hemispheres, 9 subjects, left hemisphere 14 subjects). Activation-dependent signal decrease for fQSM and increase for BOLD fMRI, respectively, could be observed for each cortical layer. fQSM showed the largest mean susceptibility decrease in Layers II/III with -1.3 ± 1.0 ppb/ -1.1 ± 1.2 ppb for M1 (-1.0 ± 1.0 ppb, when considering the left hemisphere only) and in both Layers I and II with -0.9 ± 1.0 ppb in S1 (-1.0 ± 0.9 ppb, left hemisphere only) (Figure 2b). BOLD fMRI signal increase was largest in Layer I with a maximal average increase of 3.0 ± 1.5 % in M1 (3.2 ± 1.3 %, left hemisphere only) and 2.4 ± 2.0 % in S1 (2.8 ± 2.2 %, left hemisphere only) (Figure 2d). The average number of voxels included in each layer mask ranged from 198 ± 43 voxels in Layer VI to 335 ± 36 voxels in Layer I for M1 (105 ± 16 to 170 ± 25 voxels, left hemisphere only) and from 179 ± 21 voxels in Layer Va to 262 ± 36 voxels in Layer Vb for S1 (96 ± 14 voxels to 136 ± 21 voxels, left hemisphere only) (Figure 2f) which reflects the higher number of voxels included in the outer layers of M1 due to its curvature, while numbers of voxels are more equally distributed in layers of S1 as a relatively straight section was chosen as region of interest. Supplementary Figure S1 shows the average signal evolution across subjects with standard deviations separately for each layer in M1 and S1 for both fQSM and BOLD fMRI.

**Figure 2.**
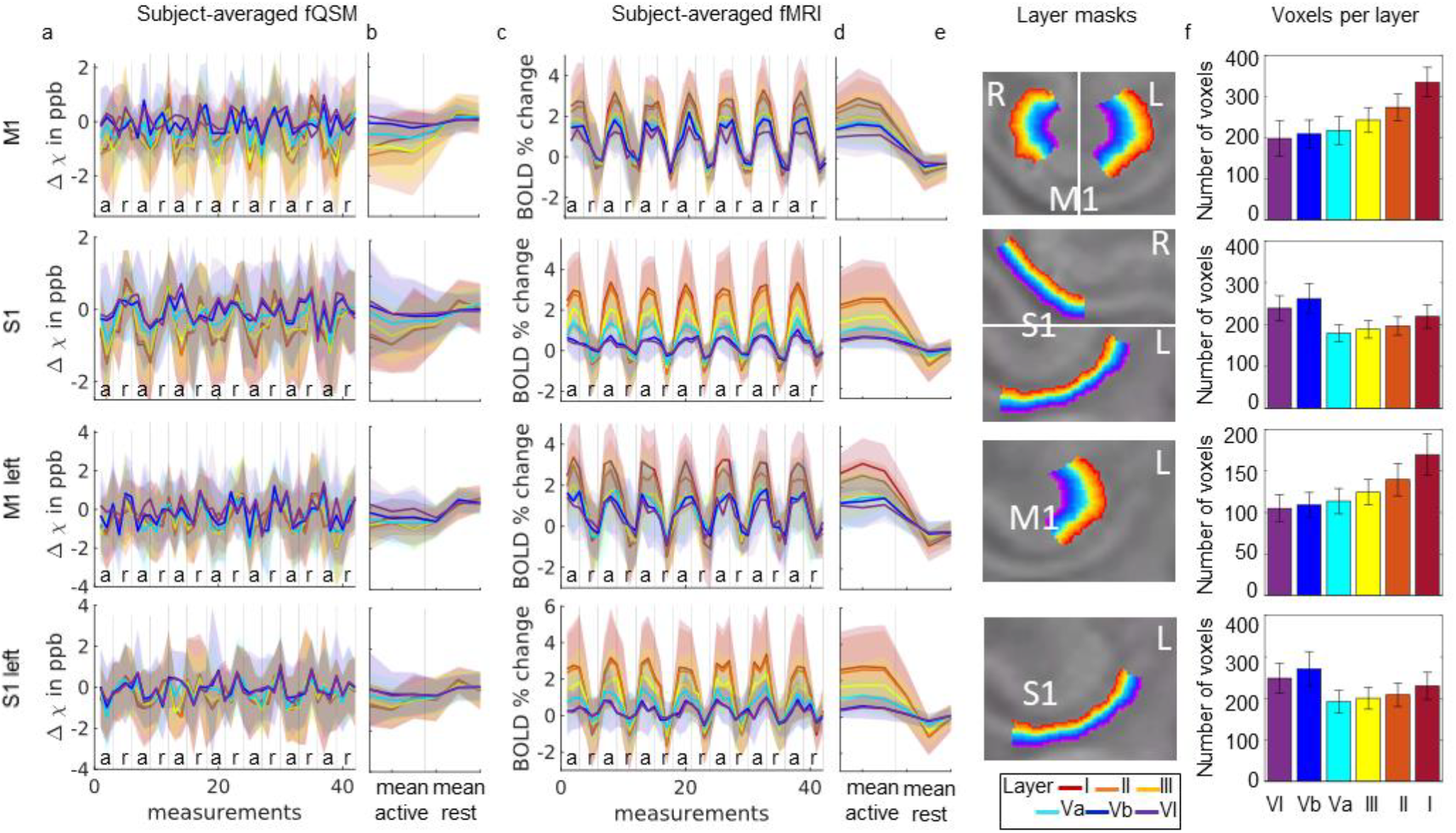
Subject-averaged fQSM and fMRI signal evolutions for Layers I (red) to VI (purple) are displayed with standard deviations as shaded curves for the motor cortex and the somatosensory cortex averaged over both hemispheres (M1, S1), and for the left motor cortex (M1 left) as well as for the left somatosensory cortex (S1 left). Subfigures b and d show time-averaged signals with standard deviations, and Subfigure e example slices of the layer segmentations. In Subfigure f, the mean number of voxels per layer across subjects is displayed with standard deviations.

When comparing activation-based signal changes for the right hemisphere between subjects who performed unilateral finger tapping with the right hand and bilateral finger tapping, the right hemisphere showed comparable results as the left hemisphere in the subjects who performed bilateral finger tapping: fQSM susceptibility decrease was largest in Layers II (-1.3 ± 2.5 ppb) and III (-1.3 ± 1.9 ppb) in M1 and Layers I (-1.3 ± 1.7 ppb) and II (-1.1 ± 1.8 ppb) in S1 (Figure 3a, b), while the percentage of BOLD fMRI signal increase was largest in Layer I (3.2 ± 1.3 % in M1, 3.4 ± 2.5 % in S1) (Figure 3c, d). For fQSM, the times series were significantly different between the subjects who performed unilateral finger tapping and the subjects who performed bilateral finger tapping for Layers I to Va (p = 0.0015 (I), p = 0.0001 (II), p = 0.0002 (III), p = 0.0409 (Va)) when assessing M1 and for Layers I (p = 0.0106), III (p = 0.0240) and Va (p = 0.0489) in case of S1. For BOLD fMRI, differences were significant for all layers for both M1 (p < 0.0001 (I), p = 0.0001 (II), p = 0.0006 (III), p = 0.0026 (Va), p = 0.0012 (Vb), p = 0.0320 (VI)) and S1 (p < 0.0001 (I), p < 0.0001 (II), p = 0.0001 (III), p = 0.0002 (Va), p = 0.0008 (Vb), p = 0.0178 (VI).

**Figure 3.**
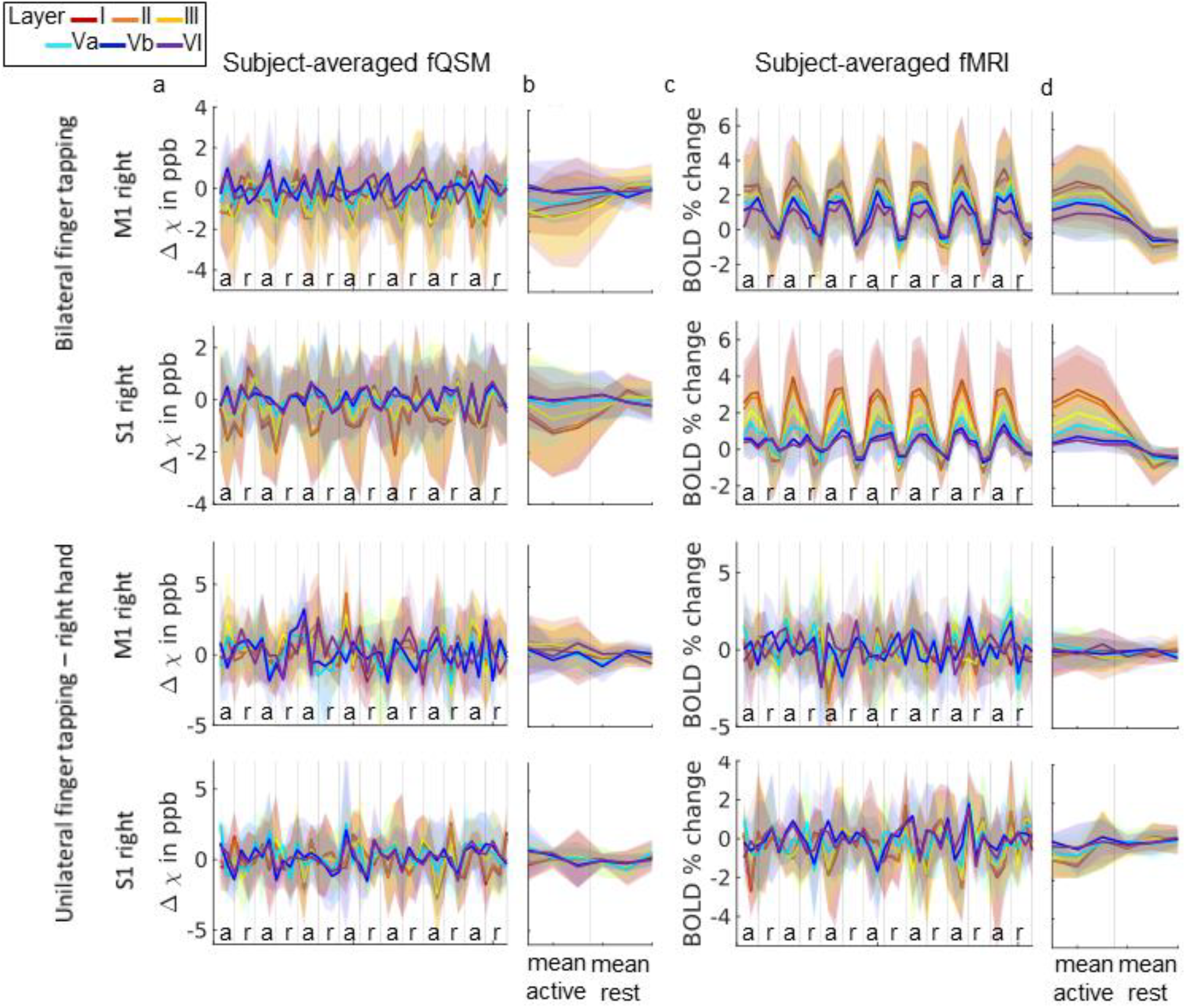
For the right hemisphere, fQSM and fMRI signal evolutions for Layers I (red) to VI (purple) are shown for M1 and S1 averaged over all subjects who performed bilateral finger tapping and unilateral finger tapping with their right hand, respectively. Standard deviations are displayed in the same color. Subfigures b and d show time-averaged signals with standard deviations.

Figure 4 illustrates the time-averaged fQSM and fMRI cortical depth profiles of M1 (left hemisphere) in each subject and corresponding maps of susceptibility and percentage of BOLD signal change. For almost all subjects, fQSM exhibited a negative peak in Layers II/ III of M1 and BOLD fMRI a peak in Layer I (in ten subjects) or layer II (in four subjects). However, for Subjects 5, 6 an almost monotone susceptibility increase was observed throughout Layers II/ III. Furthermore, double peak signals could be observed in about half of the subjects for both fQSM and fMRI, whereas the peaks for fMRI were located further towards the cortical surface compared with the negative peaks for fQSM, e.g. in Subject 4, or negative peaks for fQSM were located further towards the cortical surface within layers Vb and III/ II, e.g. in Subject 8. For Subject 2, who performed bilateral and unilateral finger tapping in two separate sessions, signal change maps and cortical depth profiles appeared highly similar regarding the location and sign of signal change as well as locations and magnitudes of negative peaks for fQSM and peaks for BOLD fMRI.

**Figure 4.**
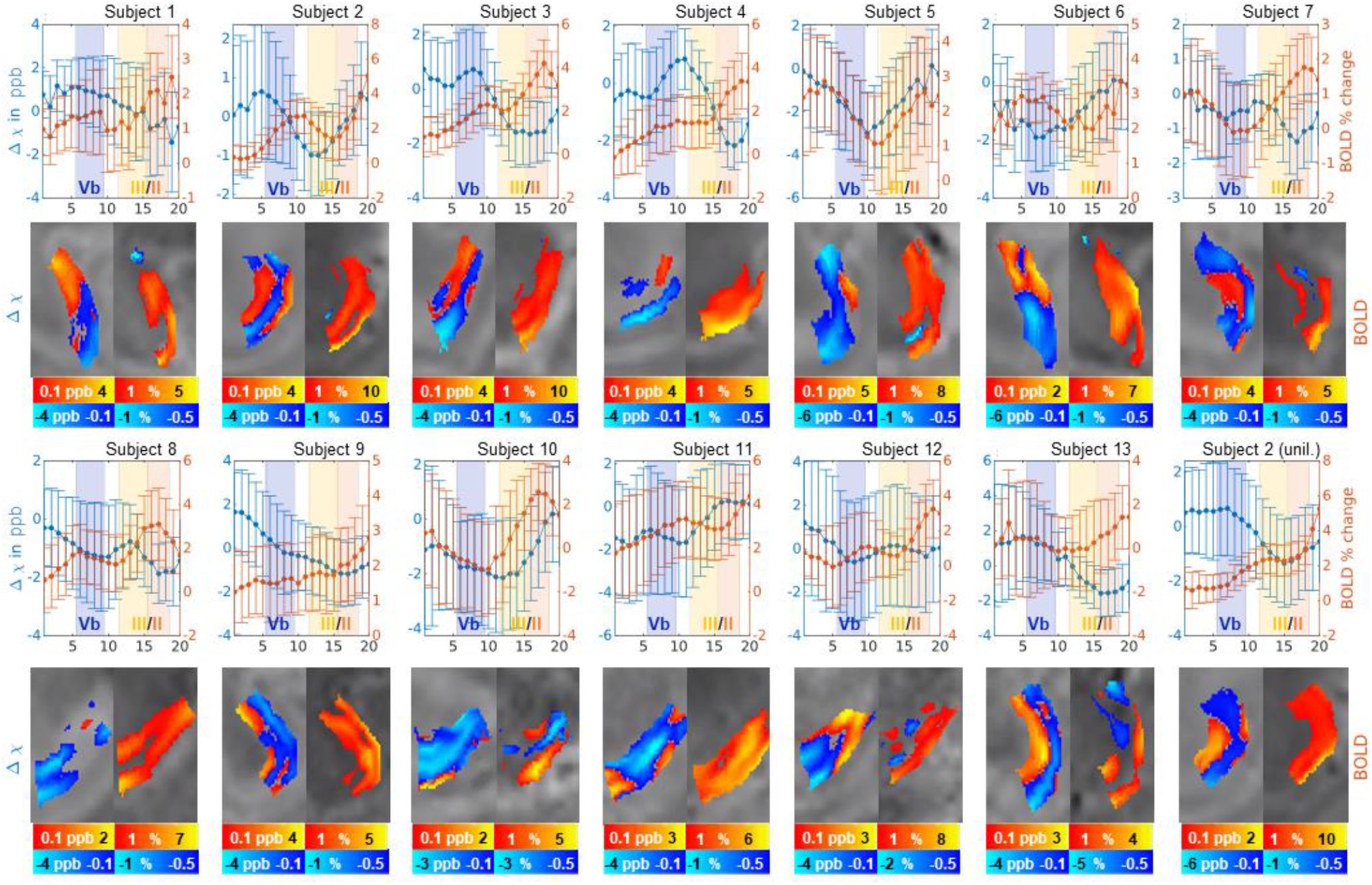
Mean (time-averaged) cortical profiles of fQSM (blue) and fMRI (orange) with standard deviations in the left M1 are shown for all 14 subjects. Corresponding susceptibility differences (in ppb) and percentage of BOLD signal change are displayed as well.

Comparing fQSM, fMRI and VASO using 3D EPI data and the data acquired with the VASO sequence in four subjects (Figure 5), fQSM computed from the BOLD acquisition of the VASO sequence as well as the BOLD fMRI part of the VASO sequence showed higher susceptibility and percent BOLD signal changes with higher standard deviations (Figure 5b, d) than the data from the 3D EPI sequence (Figure 5a, c). Mean susceptibility decrease was strongest in Layers I/ II with -3.5 ± 5.6 ppb/ -3.3 ± 6.7 ppb in M1 and in Layer I/ II with -3.0 ± 3.7 ppb/ -3.1 ± 4.2 ppb in S1, and mean percentage of BOLD signal increase was highest in Layer I in both M1 and S1 (4.6 ± 2.4 % and 2.6 ± 1.9 %). VASO (Figure 5e) showed the largest mean percent signal decrease in Layers III (-1.6 ± 4.0 %) and Va (-1.8 ± 3.6 %) in case of M1 and in Layers I/ II in S1 with -2.2 ± 0.9 %.

**Figure 5.**
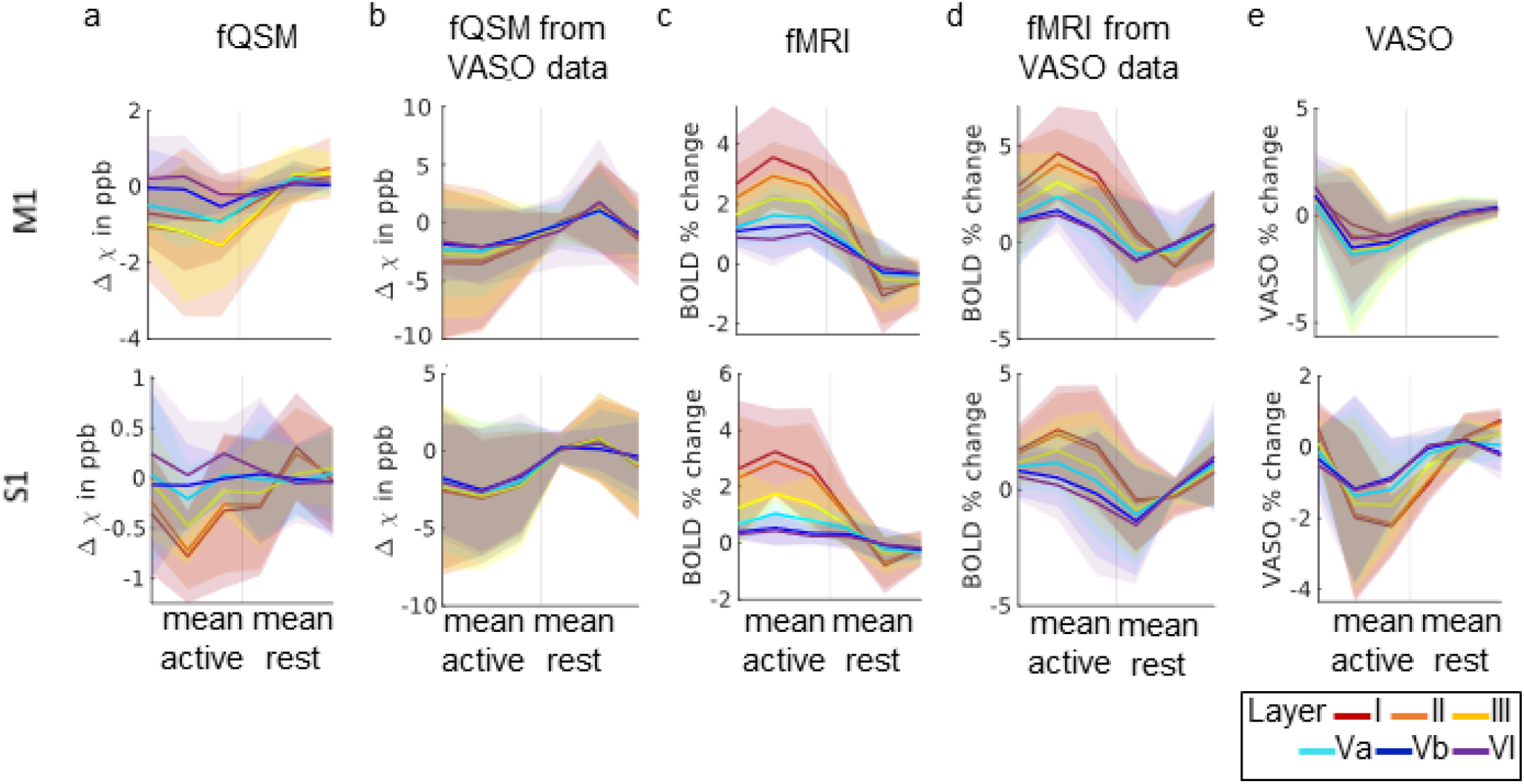
For the four subjects in which data were acquired with the 3D EPI-fQSM sequence and the VASO sequence, time-averaged fQSM, fMRI and VASO signals are shown for M1, S1 (both hemispheres averaged). fQSM signal computed from 3D EPI-fQSM data (a) as well as computed from the data of the BOLD acquisition part of the VASO sequence (b) is shown. fMRI is also displayed for both datasets, 3D EPI-fQSM (c) and the BOLD part of the VASO sequence (d). BOLD-corrected VASO data are shown in Panel e.

Both fQSM and VASO, showed positive and negative values in the z-score maps while fMRI showed predominantly positive z-scores (Figure 6a, b). Negative (positive) z-scores for fQSM and VASO were roughly colocalized, while negative z-scores in BOLD fMRI also approximately coincided with positive z-scores in fQSM/ VASO. Mean minimal negative z-scores for fQSM were significantly lower than negative z-scores for VASO or BOLD fMRI while mean maximal positive z-scores across subjects for BOLD fMRI were significantly higher than positive z-scores for fQSM and positive/ negative VASO when assessing M1. In case of S1, mean maximal positive z-scores for BOLD fMRI were significantly higher than both mean maximal positive and mean minimal negative z-scores for fQSM and VASO (Figure 6c).

**Figure 6.**
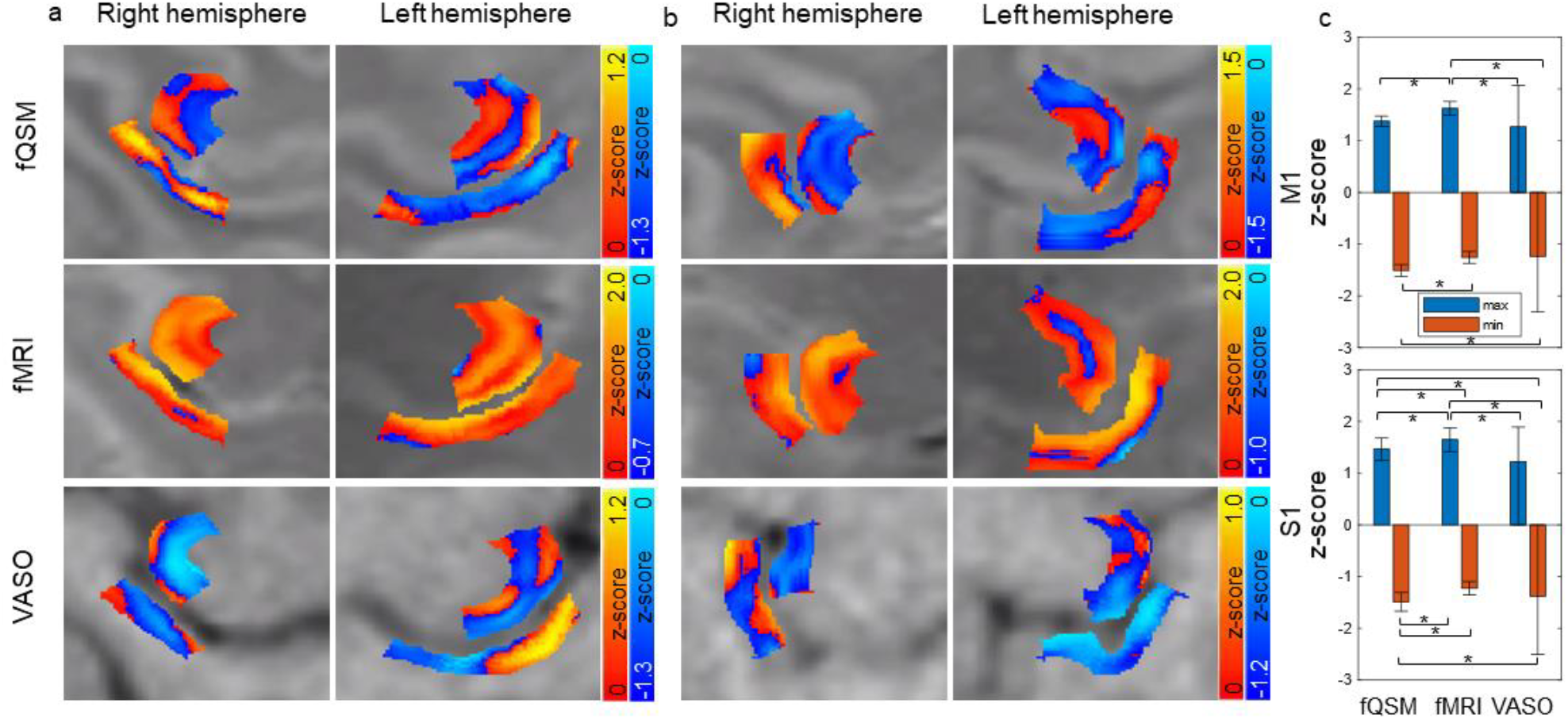
For two example subjects, in which data were acquired with both the 3D EPI-fQSM sequence and the VASO sequence, smoothed z-scores maps are displayed for fQSM, BOLD fMRI and VASO overlaid on one timepoint of the corresponding data (a, b). Panel c shows bar graphs of the mean maximal positive (blue) and mean minimal negative (orange) z-scores for fQSM, BOLD fMRI and VASO with standard deviations. Asterisks indicate statistically significant differences.

## 4. Discussion

In this study, we have demonstrated the feasibility of laminar fQSM using a 3D-EPI sequence with 0.81 mm isotropic resolution. For the investigation of M1, we observed largest activation-related susceptibility decrease in Layers II/III and in Layers I and II in S1 whereas BOLD fMRI showed the strongest signal increase in Layer I. Consistent with these findings, in cortical depth profiles fQSM showed a negative peak in Layers II/III in almost all subjects with a second negative peak in Vb in about half of the included subjects. These findings are consistent with the expected cortical depth profiles for finger tapping experiments (5,31,32). Therefore, compared to BOLD fMRI, fQSM showed less bias towards the superficial cortical layers and yielded results that were more comparable to the VASO data. It is worth noting that recent research suggests that there may be substantial individual variation across cortical profiles, which might at least in part explain the individual variations that were observed for both fQSM and fMRI (33). The magnitude of z-scores for fQSM was significantly lower than the magnitude of positive z-scores for BOLD fMRI, however, the magnitude of negative z-scores for fQSM was also significantly higher than the magnitude of negative z-scores for VASO. fQSM quantifies activation-dependent BOLD susceptibility changes, which are the basis for BOLD fMRI. Therefore, fQSM can still be expected to exhibit some bias towards the venous drainage on the cortical surface, in contrast to CBF-or CBV-based methods provided these methods are not contaminated by BOLD effect. One further advantage of laminar 3D-EPI-fQSM compared to VASO is the lower specific absorption rate and the relative ease of achieving whole-brain coverage.

Similarly to other fQSM studies (8-12), we did not only find regions of activation dependent susceptibility decrease due to increasing blood oxygenation but also areas of susceptibility increase. While the underlying mechanisms of activation dependent susceptibility increase are not fully understood and are thought to be at least partly related to the presence of larger venenous vessels which can cause artifacts in fQSM (Figure 1), positive z-scores for fQSM were roughly colocalized with positive z-scores for VASO in this study.

In the future, the use of UHF MR scanners with high-performance gradient systems (3), or higher field strength could enable even higher spatial or temporal resolutions for laminar fQSM, facilitate investigations of the fQSM hemodynamic response function and enable a better evaluation the contributions of intravascular and extravascular signal to fQSM.

Limitations of this study include the small number of subjects for which VASO data were acquired and the small number of subjects comprising different subgroups (e.g. unilateral right-hand finger tapping). We did not additionally acquire other techniques which are less sensitive to effects from venous drainage such as spin-echo BOLD fMRI or perform post-processing correction. Furthermore, the segmented layers were approximately binned into cortical Layers I to VI according to information from literature (2,34) regarding the structure of M1 and S1, which does not account for individual differences. Moreover, the phase data from the BOLD part of the VASO acquisition are not optimized for QSM, e.g. lower spatial coverage of the brain and suboptimal coil combination. When comparing, the optimized 3D-EPI fQSM data as well as BOLD fMRI and VASO, we were comparing functional activation from different acquisitions.

In conclusion, our findings demonstrate the feasibility of studying layer-dependent activation using submillimeter fQSM. Compared to BOLD fMRI, fQSM exhibits reduced bias towards venous drainage effects on the cortical surface, allowing for better localization of laminar activation.

