## Supplementary Figure S1 for "Feasibility of laminar functional quantitative susceptibility mapping"

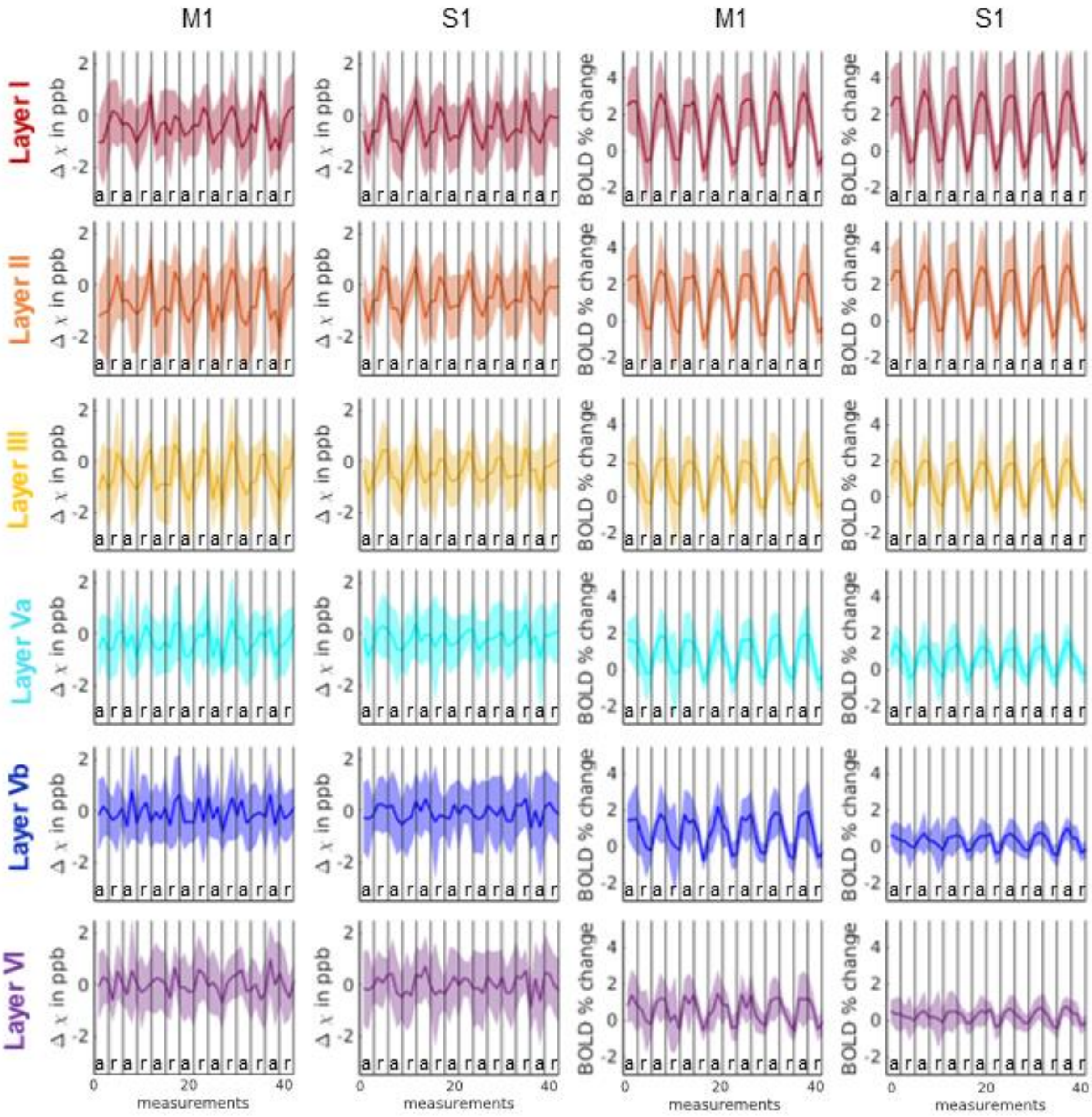


Supplementary Figure S1 shows the average signal evolution across subjects with standard deviations separately for each layer in M1 and S1 for both fQSM and BOLD fMRI. Data were average over both brain hemispheres.
